# Retention of visuo-proprioceptive recalibration in estimating hand position

**DOI:** 10.1101/2022.11.28.517441

**Authors:** Manasi Wali, Trevor Lee-Miller, Reshma Babu, Hannah J. Block

## Abstract

The brain estimates hand position using visual and proprioceptive cues, which are combined to give an integrated multisensory estimate. Spatial mismatches between cues elicit recalibration, a compensatory process where each unimodal estimate is shifted closer to the other. It is unclear how well visuo-proprioceptive recalibration is retained after mismatch exposure. Here we asked whether direct vision and/or active movement of the hand can undo visuo-proprioceptive recalibration, and whether recalibration is still evident 24 hours later. 75 participants performed two blocks of visual, proprioceptive, and combination trials, with no feedback or direct vision of the hand. In Block 1, a 70 mm visuo-proprioceptive mismatch was gradually imposed, and recalibration assessed. Block 2 tested retention. Between blocks, Groups 1-4 rested or made active movements with their directly visible or unseen hand for several minutes. Group 5 had a 24-hour gap between blocks. All five groups recalibrated both vision and proprioception in Block 1, and Groups 1-4 retained most of this recalibration in Block 2. Interestingly, Group 5 showed an offline increase in proprioceptive recalibration, but retained little visual recalibration. Our results suggested that visuo-proprioceptive recalibration is robustly retained in the short-term. In the longer term, contextual factors may affect retention.

## Introduction

There might be some limits to generalizability of our results in terms of the setup used for the experiment. We might get different results using a robotic manipulandum instead of a touchscreen as it might act as a different context. In terms of Using our hands in daily life requires the brain to interpret the available sensory signals about the position of the hand. These signals include vision, from the image of the hand on the retina, and proprioception (position sense) from sensors in the muscles of the upper limb. When different sensory cues are interpreted as arising from the same source—the hand, in this case—the brain is thought to weight and combine them to form an integrated estimate of hand position^1,2^. Estimates of hand position are used to plan appropriate motor commands to reach our goals and targets^3–5^.

Visual and proprioceptive signals about hand position are not necessarily congruent^6^, meaning that felt and actual hand position might not be the same. These mismatches vary across individuals but remain stable over time^7^. However, perturbations in the environment can cause additional incongruencies; for example, the visual estimate of the hand is shifted when the hand is immersed in a sink full of water, which refracts light. The brain compensates for such mismatches by visuo-proprioceptive recalibration, shifting the unimodal position estimates closer together to reduce the conflict^1,8–10^.

A visuo-proprioceptive mismatch also occurs during visuomotor adaptation, a process thought to involve trial-by-trial updating of the reach motor command in response to systematically perturbed visual feedback^3,11–14^. For example, cursor rotation paradigms have a visual cursor, representing the reaching hand’s position, rotated with respect to the actual hand’s movement path. This causes errors in reaching the target, which are gradually reduced as visuomotor adaptation proceeds. Proprioceptive recalibration has also been reported, with proprioceptive estimates shifting toward the rotated visual feedback^13,15,16^. Proprioceptive recalibration occurs in a variety of motor adaptation contexts, including abrupt or gradual visuomotor distortions and active or passive displacement of the hand^13,14^. Proprioceptive recalibration and visuomotor adaptation may occur in parallel but independent of each other^17^, although Tsay et al. (2022) suggest proprioceptive recalibration may in part drive visuomotor adaptation^18^. Cressman and Henriques (2010) found proprioceptive recalibration in response to a cursor rotation even in the absence of motor adaptation training trials. This study demonstrated that the cross-sensory error signal was sufficient to drive proprioceptive recalibration^16^. However, in the context of visuomotor adaptation, proprioceptive recalibration may be a response to sensory prediction errors in proprioception, in addition to the visuo-proprioceptive conflict^18^.

Changes in the motor system arising from visuomotor adaptation are retained even after the training has ended. These changes have been shown to last for several days to a year after training^19^. Retention in visuomotor adaptation is demonstrated in terms of faster relearning when exposed again to the same perturbation (savings)^20,21^ and/or movement aftereffects^22^. In a cursor rotation task, movement aftereffects would be reach errors in the opposite direction after the rotated feedback is removed. Retention of motor performance has been demonstrated in other adaptation paradigms such as velocity-dependent force field^23^ and prism adaptation^24^. Retention of proprioceptive recalibration after visuomotor adaptation training has been examined to a lesser extent, but studies have shown that proprioceptive changes after gradual visuomotor perturbations^4^ persist even after 24h. Maksimovic & Cressman (2018) found that proprioceptive recalibration was retained after 24h post-training in the form of recall, and 4 days post-training in the form of savings. They suggested that the retention of visuomotor adaptation and proprioceptive recalibration might have similar implicit processes which are linked to long term retention^3^.

It remains unclear to what degree visuo-proprioceptive recalibration in response to a visuo-proprioceptive cue conflict is retained, and what might interfere with retention. While we can speculate to some extent based on studies of proprioceptive recalibration associated with visuomotor adaptation, processes other than a response to cue conflict may be at work ^18^, and visual recalibration is generally not examined in visuomotor adaptation studies. The goal of the current study was therefore to test the retention of visuo-proprioceptive recalibration in response to a visuo-proprioceptive cue conflict, in the absence of motor adaptation. Specifically, we asked whether visuo-proprioceptive recalibration in response to a gradually imposed 70 mm mismatch was retained after participants were permitted to directly view and/or actively move their hand with no mismatch, and after 24 hours.

## Results

Various patterns of responses were observed among the participants. Generally, during Block 1, participants recalibrated both vision and proprioception to some degree. All three potential retention outcomes (Fig. 1Biii) were observed across participants. For example, the AV participant depicted in Fig. 2A recalibrated vision 43.7 mm and proprioception 29.0 mm. Early Block 2 shows similar V target undershoot but reduced P target overshoot than at the end of Block 1, suggesting this participant had full retention in vision but some forgetting in proprioception. The 24h participant depicted in Fig. 2B recalibrated vision 42.9 mm and proprioception 10.8 mm. Early Block 2 shows very little V target undershoot compared to the end of Block 1, suggesting substantial forgetting in this modality.

**Figure 1.**
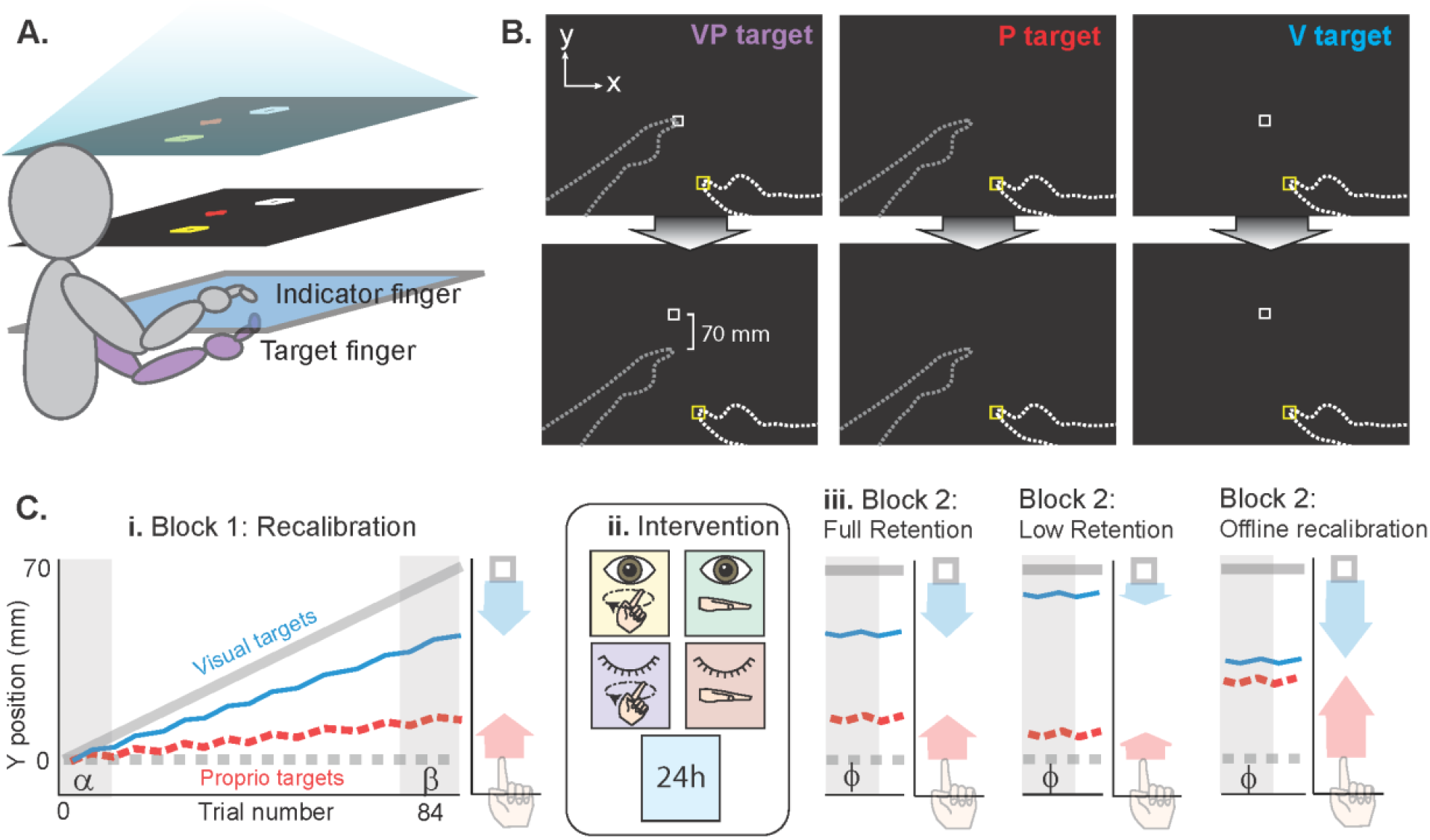
Experimental setup and Task design. **(A)** The 2D virtual reality apparatus setup. The task was viewed on the middle layer (the mirror), reflected from the projection screen above, making it appear that the display was in the plane of the bottom layer (touchscreen). **(B)** There were three target types. The bimodal VP target consisted of the target fingertip positioned beneath the touchscreen with a white box projected at the same location initially (top row), but gradually shifted forward (bottom row). The P target was the target finger with no white box, and the V target was the white box alone. The VP target was used to create the misalignment, while the unimodal targets were used to assess recalibration and retention. No direct vision of either hand was possible, and no performance feedback or knowledge of results was given. **(C)** Task design. (i) Block 1 consisted of V, P, and VP targets presented in rotating order, with a 70 mm mismatch gradually imposed in the sagittal plane (y-dimension). Blue and red lines represent estimates of the unimodal V and P targets, respectively. Colored arrows represent recalibration from early trials to late trials (grey regions). (ii) Intervention determined by random group assignment. The first four interventions consisted of a 5-minute gap between blocks; the fifth group left the lab and returned the next day for Block 2. Open eye icon: direct vision of target hand by removing mirror backing temporarily. Closed eye icon: no removal of mirror backing, no direct vision of hands. Finger on circle icon: target finger traced a circular track on the touchscreen in time with a metronome. Flat hand icon: target hand rested on the lap. (iii) Block 2 was used to assess retention of recalibration and consisted of V and P targets only. Three potential retention outcomes are illustrated schematically.

**Figure 2.**
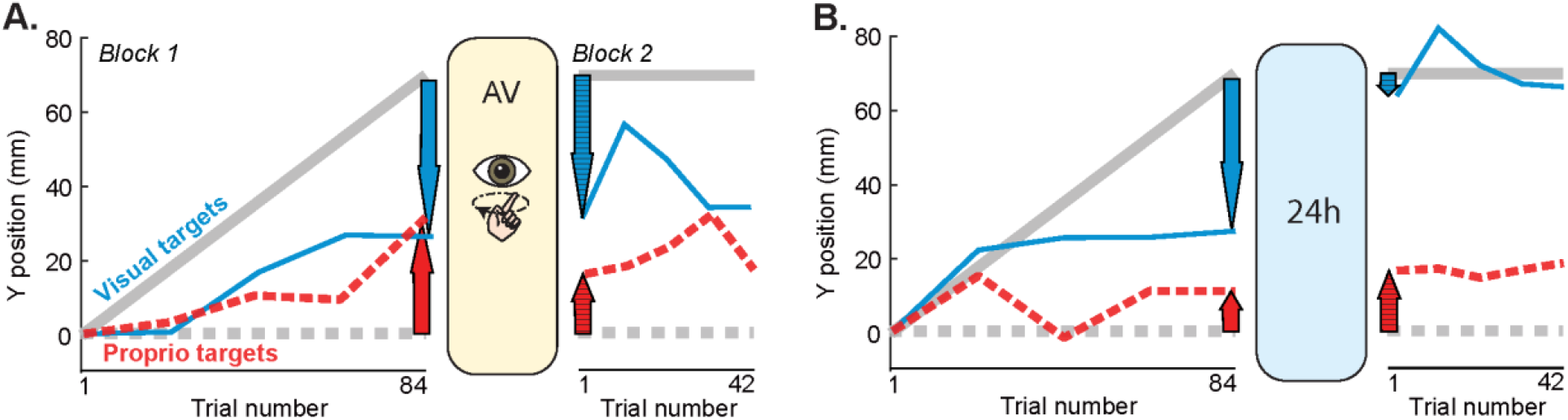
Example participants’ y-dimension indicator finger endpoints on P and V targets, averaged every 4 trials for clarity (red and blue lines). **(A)** This Active/Vision group participant recalibrated both vision and proprioception robustly during Block 1 (solid arrows). After active circle tracing with the visible target hand, the participant continued to undershoot V targets in Block 2, suggesting full visual retention. However, overshooting of P targets was reduced, which could indicate proprioceptive forgetting. **(B)** This participant in the 24H group lost most of their visual recalibration after 24 hours (crosshatched blue arrow compared to solid blue arrow), but fully retained their overshoot of proprioceptive targets.

We found that all five groups retained their recalibration, but the amount of retention varied among groups and modality (Fig. 3). Retention appears close to full (Fig. 1Biii, left) for the AV, RV, ANV, and RNV groups (Fig. 3A-D). However, the 24h group shows low retention (i.e., substantial forgetting) for vision, but offline gains for proprioception (Fig. 3E).

**Figure 3.**
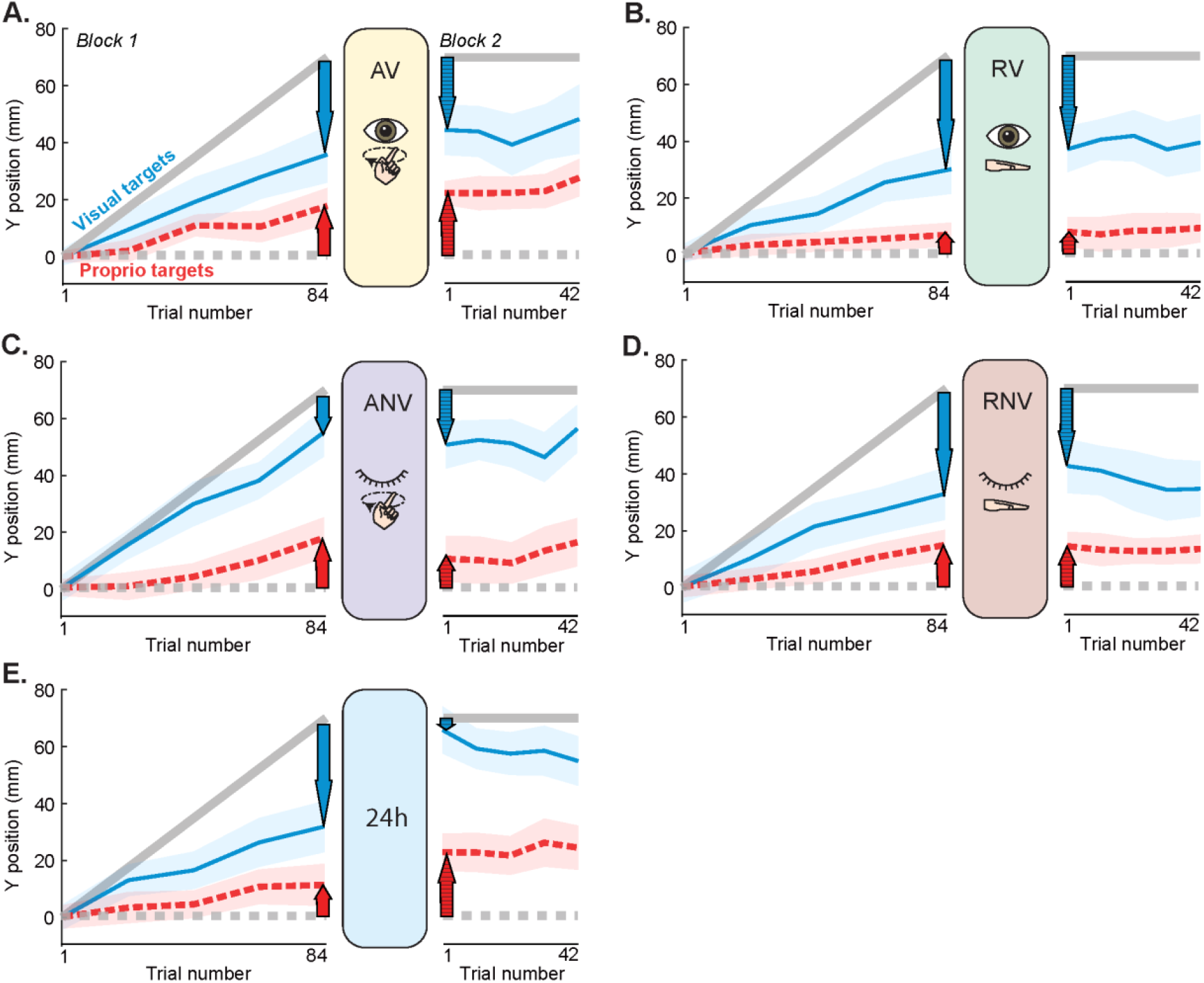
Mean task performance within groups. Blue line represents group mean estimates of V targets. Red line represents group mean estimates of P targets. Shaded regions represent SEM. **A-D**. For the active and rest, vision and no vision groups, retention (cross-hatched arrows) appears similar in magnitude to recalibration (solid arrows) for each modality, with small variations. **E**. For the 24h group, visual retention is smaller than visual recalibration, while proprioceptive retention appears larger than proprioceptive recalibration.

### Visual recalibration and retention

At a descriptive level, the AV, RV, and RNV “forgot” some of the visual change they underwent during recalibration (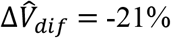-19%, and -26%, respectively; Fig. 3A, B, D). The ANV group had a small offline gain in visual recalibration (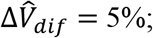; Fig. 3C), while the 24H group forgot nearly all of their visual recalibration (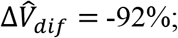; Fig. 3E). The statistical analysis for vision (mixed-model ANOVA; task stage x group) showed a significant group x task stage interaction (F_4,68_ = 7.4, *p* < 0.001, η_p_^2^ = 0.304), suggesting that groups performed the task stages differently. There was also a main effect of task stage (F_1,68_ = 24.8, p < 0.001, η_p_^2^ = 0.267) meaning that across groups, visual retention differed from visual recalibration. There was no main effect of group (F_4,68_ = 1.57, p = 0.19, η_p_^2^ = 0.084).

Post-hoc tests were conducted to determine how recalibration and retention of recalibration differed between and within the groups. This revealed that differences among the groups for visual recalibration 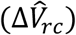 were not significant (*p > 0.5)*, meaning all the groups recalibrated vision similarly in Block 1 (Fig. 3). There were no significant group differences in raw retention of visual recalibration 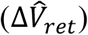, except between the RV group 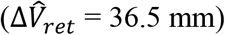 and the 24H group 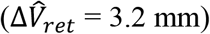.

This between-group difference of 33.3 mm was statistically significant (*p* = 0.014), meaning that the 24H group showed significantly less visual retention than the RV group (Fig. 4A). We also compared recalibration to raw retention within groups. These parameters were significantly different only for the 24H group (*p < 0.001*), meaning that the magnitude of retention 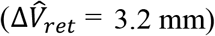 was significantly less than the amount recalibrated 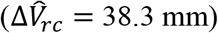 (Fig. 4A), consistent with low retention (Fig. 1Ciii).

**Figure 4.**
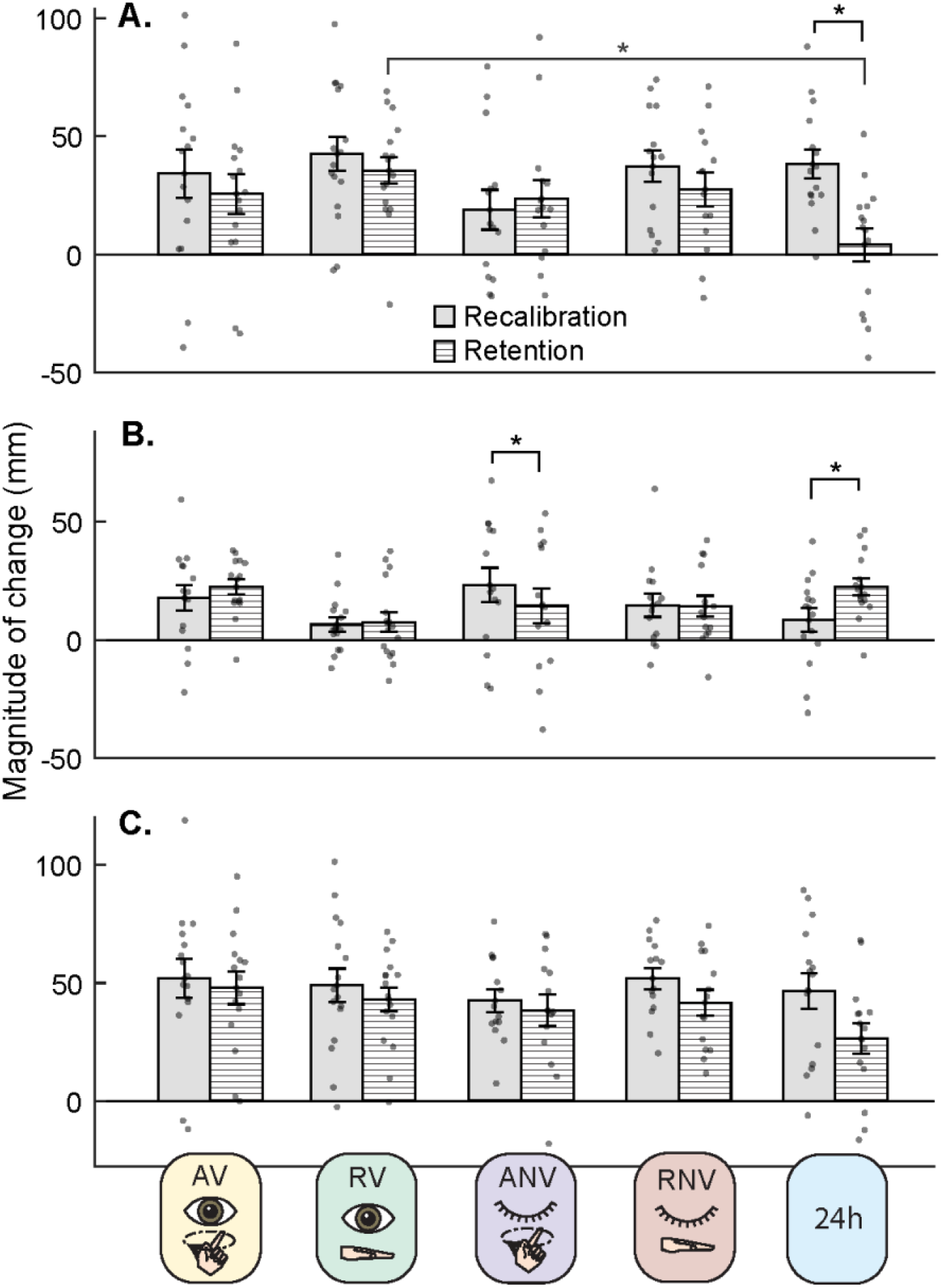
Mean group recalibration and retention with standard errors and individual data points. **(A) Vision**. There was a significant group x alignment stage interaction. Recalibration occurred similarly across groups. Retention significantly differed between Rest/Vision (RV) and 24H group. Within groups, Recalibration and Retention differed in the 24h group (35.6 mm) where retention was significantly less than recalibration. **(B) Proprioception**. There was a significant group x alignment stage interaction. Recalibration and Retention occurred similarly across groups. Within groups there was a significant difference in the recalibration and retention block for Active/No Vision and 24H group. **(C) Total**. There was no significant interaction between the group and alignment stage. *p < 0.05 for group x task stage ANOVA effects.

### Proprioceptive recalibration and retention

Descriptively, the AV, RV, and RNV groups had slight offline gains in proprioceptive recalibration (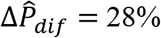, and 7%, respectively; Fig. 3A, B, D). The ANV group displayed forgetting (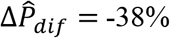, Fig. 3C), and the 24H group showed a large offline gain in proprioceptive recalibration (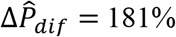, Fig. 3E). The mixed-model ANOVA (task stage vs. group) for proprioception also showed a significant group x task stage interaction (F_4,68_ = 5.88, *p* < 0.001, η_p_^2^ = 0.257), meaning that the groups behaved differently across task stages. There was no main effect of task stage (F_1,68_ = 2.23, p = 0.14, η_p_^2^ = 0.032) or group (F_4,68_ = 1.07, p = 0.38, η_p_^2^ = 0.06).

Post-hoc tests revealed that the differences among groups for proprioceptive recalibration 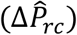 were not significant (*p > 0.5)*, meaning all the groups recalibrated proprioception similarly in Block 1 (Fig. 3). There was also no significant difference in raw proprioceptive retention 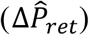 between groups (*p > 0.3*). However, we did observe significant differences between recalibration 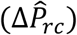 and retention 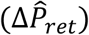 within the Active/No Vision (ANV) and the 24H group (Fig 4B). In the ANV group, 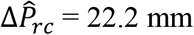 and 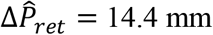 (*p = 0.024*), consistent with low retention (Fig. 1Ciii). For the 24H group, 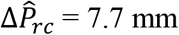 and 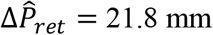 (*p < 0.001*), consistent with offline gains (Fig. 1Ciii).

We also compared raw proprioceptive retention 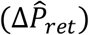 to zero in each group to ascertain whether there was any evidence that proprioceptive estimates at the beginning of Block 2 (after intervention) were different from the beginning of Block 1 (baseline). Results suggest 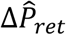 differed significantly from zero for the AV group (t_14_ = 7.26, p < 0.001, d = 1.88), the RNV group (t_13_ = 3.10, p = 0.008, d = 0.83), and the 24H group (t_14_ = 5.91, p < 0.001, d = 1.53). However, we did not find evidence of significant proprioceptive retention in the ANV group (t_13_ = 1.95, p = 0.073, d = 0.52) or the RV group (t_14_ = 1.96, p = 0.070, d = 0.51).

### Total recalibration and retention

At the group level, percent changes from total recalibration (visual plus proprioceptive) to total retention (visual plus proprioceptive), were negative, consistent with forgetting (Fig. 3). The AV group forgot the least (−4%, Fig. 3A) and the 24H group forgot the most (−46%, Fig. 3E). Statistical analysis suggests that these differences among groups are not significant. For total recalibration, there was no task stage x group interaction (F_4,68_ = 2.20, p = 0.078, η_p_^2^ = 0.12), suggesting that the groups did not perform differently across stages when visual and proprioceptive compensation is summed. Our results showed a significant main effect of task stage (F_1,68_ = 17.32, *p* < 0.001, η_p_^2^ = 0.203), meaning that within groups, total recalibration (48.9 mm) significantly differed from total retention (39.4 mm) (*p < 0.001)* (Fig. 4C). There was no main effect of group (F_4,68_ = 1.26, p = 0.30, η_p_^2^ = 0.07). One-sample t-tests indicated that the total retention was significantly different from zero for all the groups (*p < 0.001*), which is consistent with significant retention occurring in all groups, when both visual and proprioceptive retention are considered.

## Discussion

Here we investigated the retention of visual and proprioceptive recalibration in response to a 70 mm mismatch. Motor adaptation studies have suggested that even after the altered visual feedback is removed, reaching errors persist, indicating retention of the adapted motor command^21,25–27^. Similarly, studies have shown that proprioceptive recalibration that occurs during motor adaptation can be retained^3,4,28^. Our results suggest that visuo-proprioceptive recalibration is also robustly retained even after participants are given a veridical view of their actual hand and allowed to actively move the seen hand. However, the 24H group retained very little visual recalibration by the following day, while showing evidence of offline gains in proprioceptive recalibration. This indicates that the retention of visuo-proprioceptive recalibration is relatively robust in the short term, but other factors like context dependence might come into play over the longer term.

### Visual and proprioceptive recalibrations were retained to different extents

When the sum of visual and proprioceptive recalibration and retention were considered (total recalibration and retention), we found no evidence of between-group differences. All five groups showed evidence of retention. However, retention of visual recalibration and retention of proprioceptive recalibration showed distinct patterns across the groups.

In all five groups, participants began to undershoot the visual targets during Block 1, suggesting they came to feel that the visual target was closer than where it was displayed (visual recalibration). The four groups with 5-minute intervention periods all showed evidence of retention at the beginning of Block 2, meaning that they continued undershooting the visual targets. However, the 24H group showed no evidence of visual retention, meaning they “forgot” the visual recalibration that occurred during Block 1. The 24H group had much more opportunity to view their hand veridically and to use it to interact with the environment, either of which could have resulted in the forgetting we observed in the visual modality. This could involve retrograde interference of the type found in motor studies when there are competing motor memories^29^. This is outside the scope of the present study, but future work could test whether visuo-proprioceptive memories behave similarly.

It is somewhat surprising that there was no evidence of visual forgetting in the AV or RV groups, which received direct vision of their target hand. Because participants in those groups could see their actual hand (without the 70 mm visuo-proprioceptive mismatch), we expected some undoing of the visual recalibration that occurred in Block 1. Perhaps direct vision of the hand fails to undo recalibration if the hand is simply at rest, as there is nothing to require or encourage participants to use the available visual information. However, even active movement with the visible hand (AV group), where subjects presumably used their view of the hand to help guide their circle tracing, was insufficient to undo visual recalibration. One possibility is that recalibration affected a body representation that functions separately from the body representation used to control active movement. Paillard (1999) refers to these representations as the body image and the body schema, respectively, based on observations in neuropsychological patients^30^. The operation of such representations in healthy individuals is not well understood^31^, but may be a worthwhile avenue for future studies.

As with visual recalibration, in all five groups, participants began to overshoot proprioceptive targets during Block 1, suggesting they came to feel their target finger was further away than it actually was (proprioceptive recalibration). The ANV group appeared to “forget” some of their proprioceptive recalibration by Block 2, but we did not see evidence of such forgetting in the other 5-minute intervention groups. One possibility is that the active movements increased spindle activation in the periphery, making proprioception a more salient signal. Generally, a more salient signal is expected to be weighted higher and recalibrated less^1,9,32^, which could potentially translate into reduced retention of proprioceptive recalibration relative to visual. However, it is unknown whether weighting and retention of recalibration are related; we can experimentally assess weighting when visual, proprioceptive, and combined trials are present^9,33^, but here we did not include combined trials in Block 2. In any case, if active movement increased proprioceptive salience (and proprioceptive forgetting) in the ANV group, we might expect this to be true of the AV group as well. It is also possible that direct vision of the hand, in the AV group, increased salience of the visual signal. If both visual and proprioceptive signals became similarly more salient, there would be no relative difference to drive a change in weighting. Further study, perhaps including assessments of the weighting of vision vs. proprioception in Blocks 1 and 2, might clarify this question.

Interestingly, the 24H group was overshooting proprioceptive targets to an even greater extent when they returned for Block 2 the next day, consistent with an offline increase in proprioceptive recalibration. This could be considered in line with our previous work showing that magnitude of visual and proprioceptive recalibration are inversely related to each other^34^; if retention follows a similar pattern, then the increase in proprioceptive recalibration at Block 2 could be related to the reduction in visual recalibration at Block 2 for the 24H group. Another possibility is that the 24H group was influenced by contextual factors. Torres-Oviedo and Bastian (2012) showed that when learning was encoded on a treadmill (device-induced learning) with contextual information that was specific to the treadmill environment, it led to retention of context specific learning on the same apparatus (treadmill) but not in a different context (overground walking) when the errors were large^35^. In other words, proprioceptive recalibration in our study could be tied to the context of the 2D virtual reality apparatus, while visual recalibration might not be.

The 24H group differed from the other groups in important ways. Not only did more time pass before retention was tested, but participants in the 24H group also presumably spent a large portion of that time making various hand movements in various sensory contexts. Studies have shown that when visual information is available about hand position, it is frequently associated with higher spatial accuracy and relied on more heavily than proprioceptive estimates^36^. Since the 24H group participants could view their hand for a longer period and perform different activities in various workspaces, often with full view of their hand and in brightly lit conditions, they presumably spent much of that time relying heavily on their visual estimates before coming back to perform Block 2 of the task. Thus, we might conclude that not only is substantial time with direct vision of the hand required to undo recalibration, but also a modality must be relied upon (weighted heavily) to undo recalibration.

### Comparing retention of visuo-proprioceptive recalibration with retention of motor adaptation

The present study is the first, to our knowledge, to examine retention of visuo-proprioceptive recalibration that occurs purely in response to a visuo-proprioceptive mismatch. Proprioceptive recalibration can also be a response to sensory prediction errors, which occur in visuomotor adaptation paradigms^18^. Visuomotor adaptation is a form of motor learning in which movements are gradually adapted to compensate for a systematic perturbation, such as a visual cursor being rotated relative to actual hand movements^13,21^. The retention of both visuomotor adaptation and of proprioceptive recalibration in this paradigm have been investigated. While such studies do not generally consider visual recalibration, proprioceptive estimates have been observed to recalibrate concurrently with adaptation of the movements. Proprioceptive recalibration occurs with a different time course and smaller magnitude than motor adaptation^14,37^. Previous studies have reported that there are some partially shared mechanisms between motor adaptation and proprioceptive recalibration^38^. In the motor process the mechanism involves updating the forward model which uses the motor commands and the current state of body to predict the outcome of movement. It involves the sensory mechanism also as there is a mismatch between the proprioceptive limb estimates and the visual feedback cursor representing limb position^18,39^. Unlike motor learning, where learning with one effector transfers to an untrained effector^40,41^, proprioceptive changes have not been observed to transfer between hands in previous studies^42^. However, proprioceptive recalibration can generalize quite broadly in comparison to motor adaptation^43^.

Previous studies have shown that when visual feedback is removed during movement or awareness of perturbations are made more explicit^44^, the proprioceptive recalibration that accompanies visuomotor adaptation is still observed^18^. In our study, even though we purely had visuo-proprioceptive recalibration and no motor learning component, we still found that the proprioceptive and visual recalibration both were robust to task manipulations such as active movement or the veridical vision of the hand which might have led to greater awareness of the mismatch between the proprioceptive and visual estimates, similar to proprioceptive recalibration in adaptation. In addition, we have previously shown that awareness of the mismatch reduces recalibration, but only when the mismatch was larger than 70 mm^10^.

Retention of motor adaptation is usually assessed by recall (aftereffects) or savings (faster relearning). Studies have shown that retention of motor adaptation occurs even after an entire year^22^. A study by Nourouzpour et al. (2015) examined proprioceptive recalibration in the context of visuomotor adaptation, and found that after 24 h, only 46% of proprioceptive changes were retained, while 72% of motor adaptation was retained^4^. In contrast, the present study found offline gains in proprioceptive recalibration after 24 h. This is consistent with the idea that proprioceptive recalibration elicited by visuo-proprioceptive mismatch occurs by a different mechanism than proprioceptive recalibration elicited by sensory prediction error, which is present in visuomotor adaptation^18^.

Maksimovic and Cressman (2018) investigated the retention of proprioceptive recalibration associated with visuomotor adaptation over a longer duration of time (4 days). They found that the proprioceptive recalibration seemed to be recalled to some extent the next day, consistent with the other studies. But the recall was not retained after 48 h. However, they did find retention in terms of savings even after 4 days^3^. In our current study, we found that the normal vision or active movement of the hand during intervention did not lead to the return of the baseline visuo-proprioceptive calibration even after 24H which is consistent with the visuomotor adaptation studies.

### Limitations and future directions

In the present study, the greatest differences in retention were observed when comparing the four short-term groups with the 24H group, and few differences were seen among the four short-term groups. This could indicate that factors linked to the duration of time between Block 1 and Block 2 are more important than the specific activities and stimuli available during the intervention. Alternatively, it could indicate that a few minutes of intervention is insufficient to see differences between active/rest and vision/no vision, and more time is needed for the brain to recalibrate to baseline or for interference to occur. The present study cannot distinguish between these possibilities. In the future we could give the participants vision of their hands and/or perform an active task for a longer duration of time in the same workspace to keep the contextual factors similar and examine the effects on retention. It would also be useful to explore effects of visuo-proprioceptive weighting on retention of recalibration; this could yield better understanding of why proprioceptive recalibration was fully retained in the 24H group even though visual estimates were recalibrated the most. Finally, retention of visuo-proprioceptive recalibration should be assessed not only in the form of recall, but also in the form of savings, to enable better comparison with retention of visuo-motor adaptation.

Visuo-proprioceptive recalibration bears some superficial similarity to visuo-motor adaptation: it may occur trial-by-trial and acts to compensate for a perturbation. Indeed, visuo-proprioceptive recalibration may occur during visuomotor adaptation, in response to the spatial offset of the visual cursor from the hand^18^. However, in-depth study of visuo-proprioceptive recalibration in the absence of visuo-motor adaptation may clarify how these processes interact in natural behavior. As such, future research should test not only other aspects of how visuo-proprioceptive recalibration is retained, but also whether it undergoes intermanual transfer or generalizes to other tools or workspaces.

There might be some limits to generalizability of our results in terms of the setup used for the experiment. We might get different results using a robotic manipulandum instead of a touchscreen as it might act as a different context. In terms of population group, we know that older adults in the age range of 60-81 years do recalibrate similar to younger adults^45^. Therefore, our results might be generalizable to an older population group.

## Conclusions

Our study suggests that visuo-proprioceptive recalibration is robustly retained in the short term, even when the hand is viewed veridically and moved actively for several minutes. In the longer term, factors such as context dependence may play a role, and differences in retention of visual vs. proprioceptive recalibration became apparent. This study helps us provide a better insight into sensory memory contributions in visuo-proprioceptive recalibration.

## Methods

### Participants

A total of seventy-five participants (55 women, 20 men; age: 18-35, median: 21.7) were included in the study. There were 5 groups and each group consisted of 15 participants. Every participant gave written informed consent and stated that they were neurologically healthy and had normal or corrected-to-normal vision. The participants were right-handed, which was calculated using the Edinburgh handedness questionnaire^46^. All protocols were approved by the Institutional Review Board of Indiana University Bloomington.

### Recalibration task setup and targets

Participants were seated in front of a 2D virtual reality setup that consisted of a two-sided touchscreen (PQLabs), mirror, and rear projection screen, all positioned in the horizontal plane. Participants viewed the task display in the mirror, making it appear that the images were in the plane of the touchscreen (Fig. 1A). In this paradigm, the participants had to use their right index finger to indicate the position of three kinds of targets: visual (V), proprioceptive (P), and visual-proprioceptive (VP), corresponding to the sensory information available for each target (Fig. 1B). The V target was a 1 cm white square which was projected on the display whereas the P target was the left index fingertip, placed on one of the two tactile markers on the bottom touchscreen. The VP trials included both the V target and the P target. During the VP trials the subjects were explicitly told that they would be in the same location, i.e., the V component would appear directly over the P component. The participant’s right index finger was always the “indicator” finger, and the left index finger was the “target” finger on the P or VP trials only. During V trials participants were instructed to keep their target hand on their lap. The left hand was always under the touchscreen and the right indicator hand was always above. The participants did not have any direct vision of their arms or hands through the mirror, and a black disposable bib was used to obscure subjects’ view of their upper arms.

### Single trial procedure

Each trial began with the appearance of a yellow start box in one of five possible positions. The start and target positions were randomized so that the subjects could not memorize a movement direction or extent. The participants had to place their indicator finger inside the start box on the upper touchscreen for the trial to begin. Visual feedback of the indicator finger near the start box (blue circle, 8mm) was provided to help participants achieve the starting position, after which it disappeared. Participants were instructed on how to position their target hand and to keep their eyes on a red fixation cross, which appeared at random coordinates within 10 cm of the target. Finally, subjects heard a “go” signal, instructing them to begin the trial.

The participants were instructed to indicate the position of the target with as much accuracy as possible at their own pace (no speed restrictions). They were instructed to lift their indicator finger off the glass and place it down at the perceived location of the target and not to drag it on the touchscreen. When satisfied with their indicator finger’s position, subjects were asked to hold their position for 2s for the system to register their touch. A camera-click sound indicated trial completion. No performance feedback was provided; thus, participants had no way of knowing how accurately they were indicating target positions.

### Session design

There were 5 groups in total and every group had to perform 126 trials in two blocks, after first performing 8 practice trials. Block 1 was used to create a visuo-proprioceptive mismatch in target finger perception and measure participants’ recalibration of vision and proprioception in response. Block 2 was used to test the retention of visual and proprioceptive recalibration when the visuo-proprioceptive mismatch was no longer explicitly presented (Fig. 1C).

Block 1 (84 trials) consisted of 21 V trials, 21 P trials and 42 VP trials, presented in the repeating order VP-V-VP-P. A 70 mm misalignment was gradually imposed in Block 1 between the visual and proprioceptive components of target position. This was done by shifting the V component away from the P component of VP targets, 1.67 mm at a time. The V component gradually shifted away from the P component in the forward direction (i.e., positive y direction), such that by the end of the misalignment block, the V component was 70 mm further away from the subject than the P component (Fig. 1B). On V target trials, the V target was projected at the location of the most recent VP trial.

After Block 1, Groups 1-4 had a 5-minute intervention period before beginning Block 2 (Fig. 1C). The intervention consisted of either one minute of active movement of the target (left) hand or rest, with or without direct visual feedback of the target hand. Group 1 was the active – vision (AV) group, Group 2 was rest – vision (RV), Group 3 was active – no vision (ANV), Group 4 was rest – no vision (RNV) Fig. 2(ii). During the intervention, direct visual feedback was achieved by removing the foam board that is normally positioned under the mirror, which is half-silvered. A lamp beneath the mirror makes the hands and lap clearly visible to the subject when this foam board is removed. The active groups performed a circle-tracing task for one minute, using their target finger to trace around a circular plastic plate which was placed at the same position as the tactile markers on the bottom touchscreen. Rest groups kept their hands on their laps for the entire 5-minute period with or without vision, and active groups did so for the 4 minutes they were not circle-tracing. A metronome was used to pace the circle tracing task in the active groups. The subjects had to complete one circle with every tone per second for a minute. To make it uniform across subjects the rest groups also had to listen to the metronome for the first 60 seconds. Group 5 (24h) participants left the lab at the end of Block 1 and returned the following day to complete Block 2. Group 1-4 participants began Block 2 immediately following the 5-minute intervention period.

Block 2 (42 trials) consisted of 21 V and 21 P trials presented in alternating order, with no VP trials. In Block 2, the V target remained where it was at the end of Block 1 (70 mm forward from the P target) and was used to determine the retention of visual recalibration from Block 1. The P trials were used to determine the retention of proprioceptive recalibration from Block 1. After Block 2, participants were asked whether they had perceived the white box to be offset from their target finger at any point. If they said yes, they were asked to report the direction of perceived offset and to estimate the spatial magnitude. 20 subjects in total reported perceiving some amount of forward offset: 2, 5, 6, 2 and 5 respectively in groups 1 – 5.

### Data Processing

We analyzed data consisting of indicator finger endpoints on V and P targets early and late in Block 1 and early in Block 2. Because the visuo-proprioceptive mismatch occurred only in the sagittal plane (y-dimension), our analyses considered only this dimension of the indicator finger endpoints. Individual data were processed and analyzed using MATLAB version 2019b (MathWorks). Statistical analyses were performed with SPSS version 28. The data is publicly available at https://osf.io/n4tbj/.

#### Recalibration

Proprioceptive recalibration 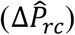 was calculated by finding the difference between the means of the last four 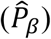 and first four 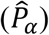 P trial endpoints in Block 1 (Fig. 1Ci):

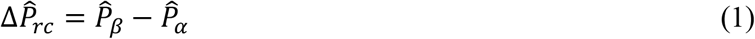

Visual recalibration 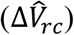 was calculated as the difference between the means of the last four 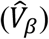 and first four 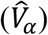 V trial endpoints in Block 1 (Fig. 1Ci), subtracted from 70 to account for the change in V target position between these timepoints.

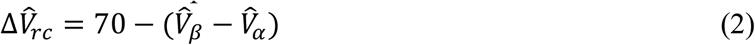

#### Retention

Raw proprioceptive retention 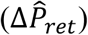 was calculated as the difference between the mean of the first four P trial endpoints in Block 2 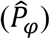 and the mean of the first four P trial endpoints of Block 1 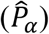 (Fig. 1Ci and iii):

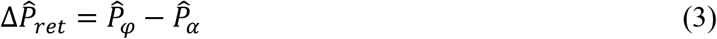

Raw visual retention 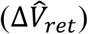 was calculated as the difference between the mean of the first four V trial endpoints of Block 2 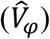 and the mean of the first four V trial endpoints in Block 1(Fig. 1Ci and iii), subtracted from 70 to account for the change in V target position between these timepoints.

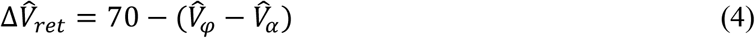

#### Percentage difference in retention

For descriptive purposes, we also computed the difference between recalibration and raw retention for each unimodal target 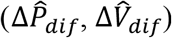:

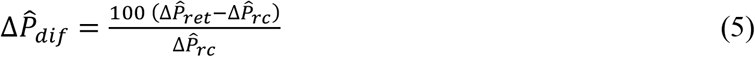

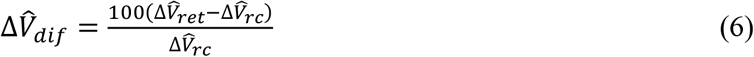

This value reflects how much retention was evident in Block 2, given how much recalibration occurred in Block 1. In other words, one participant might recalibrate proprioception 20 mm and show retention of 10 mm, meaning he “forgot” half of his recalibration, while another person might recalibrate proprioception only 10 mm but show 10 mm of retention, meaning he retained all of his recalibration. These two individuals would have equal raw retention, but the retention difference of the first person would be -50% (forgot half) versus 0% for the second person (forgot nothing).

### Statistical Analysis

To determine the effect of group intervention on recalibration and retention of recalibration, we performed a mixed model ANOVA with task stage (recalibration, raw retention) as the within-subjects factor, and group (AV, RV, ANV, RNV, 24h) as the between-subjects factor for each of vision, proprioception, and total (sum of visual and proprioceptive values). Shapiro-Wilk’s tests revealed that the group data were normally distributed. Effect sizes were computed using partial eta squared, η_p_^2^ values. Post-hoc tests with Bonferroni adjustments were performed on significant interactions. Significance was considered at an alpha level of .05. To further examine the effect of intervention on retention, we performed a one-sample *t* test on each group’s raw retention against a test value of 0. Effect sizes were computed using Cohen’s d. We excluded one subject each from group 3 and 4 due to technical issues with the touchscreen during the data collection.

## CRediT authorship contributions

MW: Conceptualization, Data curation, Formal analysis, Investigation, Methodology, Software, Visualization, Writing – original draft. TLM: Formal analysis, Writing – review & editing. RB: Conceptualization, Writing – review & editing. HJB: Conceptualization, Formal analysis, Funding acquisition, Supervision, Writing – review & editing.

## Data availability statement

The dataset generated during and/or analyzed during the current study are available in the OSF repository at https://osf.io/n4tbj/.

